## Supplementary material for "Linear discriminant analysis reveals hidden patterns in NMR chemical shifts of intrinsically disordered proteins": LDA vs. other classification methods

### 1 LDA vs. other classification methods

In this section we compare the performance of LDA with that of Quadratic Discriminant Analysis (QDA), K-Nearest Neighbours (KNN) and Support Vector Machines (SVM). The same 18 proteins from the BMRB described in the main text were used for leave-one-out cross-validation of all mentioned methods (Fig. S1). KNN was implemented with votes from 9 nearest neighbours, *taxicab* metric to measure distance ($L_1$ norm) and using the inverse of the distance as weighting function for the votes. The SVM model was constructed using a one-versus-all coding design (for each binary learner, one class is positive and the rest are negative) and was implemented using polynomials of grade 3 as kernel function to map the data onto higher dimensions.

We further analyzed the classification performance of each method by computing their sensitivity and specificity for each amino acid residue type (Fig. S2), where:

$$Sensitivity = \frac{TruePositives}{TruePositives + FalseNegatives}$$
$$Specificity = \frac{TrueNegatives}{TrueNegatives + FalsePositives}$$

For a given amino acid type (e.g. alanine), True Positives are accurately classified residues of the given type (alanine classified as alanine) and False Positives denotes residues from other amino acid types classified into it (serine classified as alanine). Similarly, True Negatives denotes residues from other amino acid types not classified into the given type (serine classified as something else than alanine) and False Negatives are wrongly classified residues of the given type (alanine classified as serine). Sensitivity measures the true positive rate, or the ability of a method to correctly classify residues of a given amino acid type, while specificity measures the true negative rate, or the ability of a method to correctly disregard residues that are not of a given amino acid type [?]. The code for these tests was written in Matlab R2021a using the Statistics and Machine Learning toolbox.

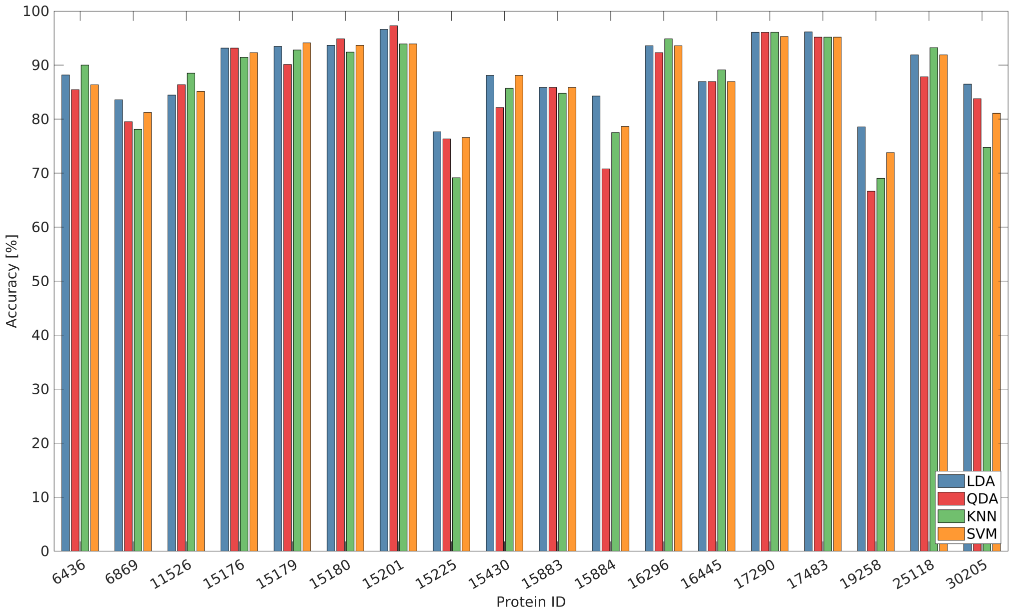


**Fig 1.** Results from leave-one-out cross-validation for various classification methods using the 18 proteins from BMRB and chemical shifts subset (iii): ($H^N$, N, CO, $C_\alpha$, $C_\beta$, $H_\alpha$ and $H_\beta$).

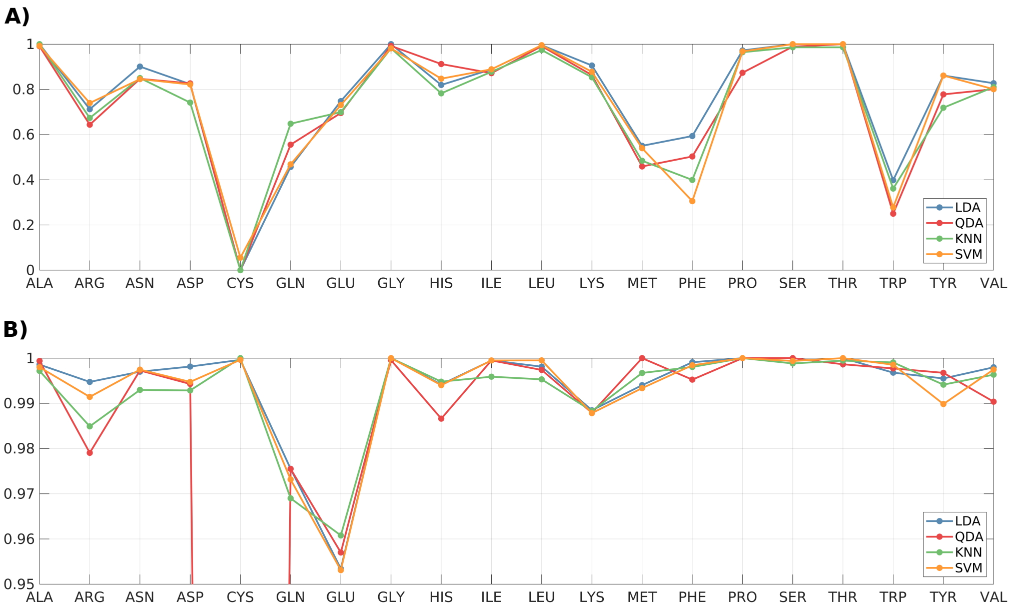


**Fig 2.** Comparison of classification performance for various methods. Results obtained from performing leave-one-out cross-validation using the 18 proteins from the training set and chemical shifts subset (iii). A) Sensitivity and B) specificity values for each amino acid type. All methods show a low performance for cysteine since there are only 5 residues of this type in the training set. In particular, this class needs to be removed for QDA since there are fewer samples than variables (which makes the class covariance matrix for this residue type singular).

A summary of the results obtained for LDA and all other classification methods is shown in Table S1. LDA is superior in every aspect. And although performance values

| Parameter | LDA | QDA | KNN | SVM |
|---|---|---|---|---|
| mean accuracy [%] | 89.43 | 87.25 | 87.29 | 88.11 |
| variance | 34.05 | 74.40 | 79.88 | 46.93 |
| mean sensitivity | 0.773 | 0.742 | 0.740 | 0.750 |
| mean specificity | 0.994 | 0.943 | 0.993 | 0.993 |

**Table 1.** Summary of classification performances. Mean accuracies are weighted by the number of residues in the test protein. Variances were computed from the values displayed in Fig. S1. Mean sensitivities and specificities were computed from the values shown in Fig. S2

for other methods may come close to that of LDA, what makes LDA come through as the best choice for protein mapping is its reduced variance on classification accuracy. In short, LDA is the best method to make consistent classification predictions across different IDPs.
